# The dual role of TonB genes in turnerbactin uptake and carbohydrate utilization in the shipworm symbiont *Teredinibacter turnerae*

**DOI:** 10.1101/2023.02.23.529781

**Authors:** Hiroaki Naka, Margo G. Haygood

**Affiliations:** Department of Medicinal Chemistry, the University of Utah; Division of Genetics, Oregon National Primate Research Center, Oregon Health & Science University

## Abstract

*Teredinibacter turnerae* is an intracellular bacterial symbiont that resides in the gills of shipworms, wood-eating bivalve mollusks. This bacterium produces a catechol siderophore, turnerbactin, required for the survival of this bacterium under iron limiting conditions. The turnerbactin biosynthetic genes are contained in one of the secondary metabolite clusters conserved among *T. turnerae* strains. However, Fe(III)-turnerbactin uptake mechanisms are largely unknown. Here, we show that the first gene of the cluster, *fttA* a homologue of Fe(III)-siderophore TonB-dependent outer membrane receptor (TBDR) genes is indispensable for iron uptake via the endogenous siderophore, turnerbactin, as well as by an exogenous siderophore, amphi-enterobactin, ubiquitously produced by marine vibrios. Furthermore, three TonB clusters containing four *tonB* genes were identified, and two of these genes, *tonB1b* and *tonB2*, functioned not only for iron transport but also for carbohydrate utilization when cellulose was a sole carbon source. Gene expression analysis revealed that none of the *tonB* genes and other genes in those clusters were clearly regulated by iron concentration while turnerbactin biosynthesis and uptake genes were up-regulated under iron limiting conditions, highlighting the importance of *tonB* genes even in iron rich conditions, possibly for utilization of carbohydrates derived from cellulose.

## Introduction

Iron is an essential nutrient for almost all living organisms including bacteria. However, available free iron is extremely limited in the marine environment due to its insolubility in the presence of oxygen, and in the host due to iron chelation by host iron-binding proteins; thus the amount of available free iron is much lower than the amount bacteria require for their proliferation. Therefore, bacteria have evolved active transport systems to sequester sufficient amounts of iron to survive and prosper in those environments (Crichton, 2016). One of these systems is siderophore-mediated iron transport. Siderophores are small molecule iron-chelating compounds synthesized by a nonribosomal peptide synthetase (NRPS) system or NRPS-independent pathway. Siderophores exported to external environments form stable complexes with ferric iron, and in Gram-negative bacteria, Fe^3+^-siderophore complexes are transported to the bacterial cytosol via specific outer membrane receptors across the outer membrane, and ABC or MSF type siderophore transporters across inner membranes (Crosa and Walsh, 2002;Cuiv et al., 2004;Raymond and Dertz, 2004;Winkelmann, 2004;Hannauer et al., 2010;Reimmann, 2012). Gram-negative bacteria require TonB complexes typically composed of TonB, ExbB and ExbD, that locate in the inner membrane, to transduce energy derived from proton motive force to the Fe^3+^-siderophore specific outer membrane receptors for their activity (Braun, 1995;Postle and Kadner, 2003;Noinaj et al., 2010). Although essential, an excess amount of iron is toxic due to its radical potential, therefore the expression of genes required for iron transport are tightly regulated by the concentration of iron to maintain a suitable cellular iron concentration (Andrews et al., 2003). It has also been demonstrated in many bacteria that iron influences not only the expression of iron metabolism genes but also acts as a signal that regulates the expression of genes that affect bacterial adaptation to environmental and/or host conditions (Crosa, 1997;Andrews et al., 2003;Fleischhacker and Kiley, 2011).

Shipworms of the family Teredinidae are marine bivalve mollusks most of which bore wood and consume wood as a nutrient source (Turner, 1966;Distel et al., 2011). To utilize wood as a nutrient, insoluble lignocellulose needs to be broken down into soluble forms of carbohydrate. This enzymatic activity relies on symbiotic gammaproteobacteria that reside in bacteriocytes in the gills (Waterbury et al., 1983;Distel et al., 2002a;Luyten et al., 2006;Ekborg et al., 2007). *Teredinibacter turnerae* is the first bacterial symbiont isolated from shipworms. This bacterium produces cellulolytic enzymes and fixes atmospheric nitrogen that could contribute to shipworm metabolism in woody environments where the amount of nitrogen is restricted (Distel et al., 2002b;Lechene et al., 2007;Altamia et al., 2014;O’Connor et al., 2014). *T. turnerae* T7901 carries many secondary metabolite gene clusters and production of bioactive compounds has been reported (Elshahawi et al., 2013;Han et al., 2013;Miller et al., 2021;Miller et al., 2022). One of the secondary metabolite gene clusters, Region 7, carries the genes that are responsible for the biosynthesis of siderophore turnerbactin (Han et al., 2013). Sequencing and metagenomic analysis revealed that the Region 7 cluster and its relatives were found to fall within the gene cluster family GCF_8, members of which occur in all *T. turnerae* strains sequenced as well as other shipworm symbiotic bacteria, indicating the importance of this cluster for the physiology of shipworm symbiotic bacteria (Altamia et al., 2020). The *tnbF* gene encoding a non-ribosomal peptide synthetase in this cluster was shown to be essential for the biosynthesis of turnerbactin and survival of this bacterium under iron limiting conditions (Han et al., 2013). Turnerbactin was detected in the shipworm, *Lyrodus pedicellatus*, harboring *T. turnerae*, suggesting the potential importance of turnerbactin in the symbiotic state. *T. turnerae* might have elevated iron requirements due to the need to synthesize iron rich nitrogenase (Han et al., 2013). It has been reported that *T. turnerae* carries two TonB gene clusters, TonB2 and TonB3, that resemble clusters found in marine vibrios although the function of those genes has yet to be characterized (Kuehl and Crosa, 2010). In this work we show the essential role of the *fttA* gene encoding the Fe^3+^-turnerbactin outer membrane receptor for iron acquisition in *T. turnerae*. Additionally, two of four *tonB* genes in the genome were indispensable for growth under iron limiting conditions. These *tonB* genes were further found to be necessary for the efficient growth of *T. turnerae* when cellulose was used as a sole carbon source. Furthermore, we report that *tonB* genes in *T. turnerae* T7901 are not clearly regulated by iron as compared with other iron transport-related genes, suggesting that *T. turnerae* requires TonB genes even under iron rich condition to utilize carbohydrate(s) originating from cellulose.

## Materials and Methods

### Strains, plasmids and growth media

Bacterial strains and plasmids used in this study are listed in **Table S1** while PCR primers are listed in **Table S2**. *Teredinibacter turnerae* strains were cultured at 30 °C in a modified chemically-defined shipworm basal medium (SBM) containing NaCl (17.94 gm/L), NH_4_Cl (250 mg/L), Na_2_SO_4_ (3.01 gm/L), NaHCO_3_ (0.147 gm/L), Na_2_CO_3_ (10.5 mg/L), KCl (0.5 gm/L), KBr (73.5 mg/L), H_3_BO_3_ (22.36 mg/L), SrCl_2_·6H_2_O (18 mg/L), KH_2_PO_4_ (15.24 mg/L), C_6_H_8_O_7_ (2.75 mg/L), NaF (2.25 mg/L), Na_2_MoO_4_·2H_2_O (2.4 mg/L), MnCl_2_·4H_2_O (1.81 mg/L), ZnSO_4_·7H_2_O (0.22 mg/L), CuSO_4_·5H_2_O (0.079 mg/L), Co(NO_3_)_2_·6H_2_O (0.049 mg/L), HEPES (4.77 gm/L, pH = 8.0), and appropriate amounts of carbon sources. MgCl_2_·6H_2_O, CaCl_2_·2H_2_O and ferric ammonium citrate (FAC) were supplemented in the medium. Sucrose (0.5 %), cellulose (sigmacell 101)(0.2 %), and carboxymethylcellulose (0.5 %) were used as carbon sources, and agar (1 %) was added to prepare solid media. Under our standard growth conditions, which include 50 μM of MgCl_2_·6H_2_O and 10 μM of CaCl_2_·2H_2_O, cell aggregation was observed. However, we found that by a reducing the concentration of MgCl_2_·6H_2_O (0.05 μM) and CaCl_2_·2H_2_O (0.5 μM) in the SBM medium, *T. turnerae* grew without aggregation. *E. coli* strains were cultured in LB broth or agar. Thymidine at 0.3 mM (f/c) was supplemented for the growth of *E. coli* π3813. When required, antibiotics were supplemented in the growth medium at the following concentration: ampicillin (Amp) at 100 μg/ml for *E. coli*, kanamycin (Km) at 50 μg/ml for *E. coli* and *T. turnerae* and carbenicillin (Carb) at 100 μg/ml for *T. turnerae*.

### Construction of plasmids

The plasmid pHN31(pDM4-Km), used for mutant construction was constructed as follows. The kanamycin resistance cassette from pBBR1MCS-2 (Kovach et al., 1995) was PCR-amplified using Km-F-*EcoR*V and Km-R-*EcoR*V primers, and ligated into T-vectors. After confirming the nucleotide sequences, the Km cassette was cloned into the *EcoRV* site of pDM4 (Milton et al., 1996), generating pHN31.

To express genes in *T. turnerae*, we used the pHN33(pPROBE-tacP-GenP) plasmid constructed as follows. pMMB208 was digested with *Sca*I and *AgeI*, and the DNA fragment containing *lac*I, the *tac* promoter and a multiple cloning site was ligated into the corresponding restriction enzyme sites of pPROBE’-gfp[ASV] (Miller et al., 2000), generating pHN32. The DNA fragment containing the gentamicin resistance gene promoter from pBBR1MCS-3 (Kovach et al., 1995) were PCR-amplified using primers, GenP-F-*Hind*III and GenP-R-*Sal*I, and cloned into T-vector. After confirmation of the nucleotide sequence, the plasmid was digested by *Hind*III and *Sal*I, and the promoter sequence was ligated in the corresponding restriction enzyme sites of pHN32 plasmid, generating pHN33. pPROBE-gfp[ASV] was a gift from Steven Lindow (Addgene plasmid # 40166; http://n2t.net/addgene:40166; RRID:Addgene_40166)

### Mutant construction and complementation

DNA fragments of upstream and downstream regions of the target genes to be mutated were combined by splicing by overhang extension PCR with modification as described before (Senanayake and Brian, 1995;Naka et al., 2018), and the PCR-amplified fragments were ligated into pGEM-T easy (Promega). After sequence confirmation, the deletion fragments were ligated into the corresponding restriction enzyme sites of pHN31. The plasmids thus obtained were transformed into *E. coli* strains S17-1λpir or π3813 (thymidine auxotroph), and conjugated into *T. turnerae* T7901. When *E. coli* π3813 was used, thymidine (f/c 0.3 mM) was supplemented to the growth medium, and *E. coli* π3813 that carries pEVS104 (Stabb and Ruby, 2002) was used as a conjugation helper strain. To counterselect *E. coli*, 1st recombinants were selected by plating exconjugants on SBM-N-cellulose plates (for S17-1λpir conjugation) with Km (50 μg/ml) or SBM-N-sucrose without thymidine plates (for π3813 conjugation) supplemented with Km (50 μg/ml). 1st recombinants thus obtained were grown in liquid medium without antibiotics, streaked on SBM-N containing 15% sucrose, and incubated until colonies were formed. The deletion mutants were obtained by screening the colonies that were sensitive to Km, by colony PCR using primers. To complement mutants, DNA fragments that contain wild type genes and their potential ribosomal binding sites were PCR-amplified, and cloned into T-vectors. After sequence confirmation, the DNA fragments were cloned into pHN33, and the plasmid was conjugated into *T. turnerae* as described above.

### RNA extraction

All glassware was soaked in a 10% hydrochloric acid bath then rinsed with milliQ water, to remove iron. *T. turnerae* T7901 and its derivatives were grown in iron limiting (L-SBM-N-sucrose with 0.1 μM FAC) and iron rich (L-SBM-N-sucrose with 10 μM FAC) until exponential phase (OD600 0.2-0.3), and cell pellets were resuspended in TRIzol Reagent (Invitrogen), and the samples were kept in a −80 °C freezer until processed. Total RNAs were extracted by the Trizol-RNeasy hybrid protocol (Lopez and Bohuski, 2007). During RNA extraction, contaminated DNA was digested by 3 times treatment with RNase-Free DNase Set (Qiagen), and the absence of DNA contamination in extracted RNA was confirmed by PCR.

### Quantitative RT-PCR

cDNA was synthesized from total RNA (1 μg) as a template using Superscript III Reverse Transcriptase and random hexamer primers (Invitrogen), and quantitative PCR were performed by StepOnePlus™ Fast Real-Time PCR System (Applied Biosystems) using Power SYBR™ Green PCR Master Mix (Applied Biosystems). The fold change of gene expression in two different conditions was measured by calculating ΔΔCt values as described in (Livak and Schmittgen, 2001).

## Results

### Characterization of the Fe-turnerbactin outer membrane receptor gene, *fttA*

One of the secondary metabolite clusters, Region 7 of *T. turnerae* T7901 contains nonribosomal peptide synthetase (NRPS) genes (**Figure 1)**, and the major NRPS gene, *tnbF* was shown to be essential for the siderophore turnerbactin production (Han et al., 2013). The first gene of Region 7, *TERTU_RS18025* (old locus tag, *TERTU_4055*) was annotated as a homologue of the TonB dependent outer membrane receptor (TBDR)g gene, CCD03052, from *Azospirillum brasilense* Sp245 (Han et al., 2013). Further comparison of the predicted amino acid sequence of *TERTU_RS18025* (named *fttA* in this study) with known Fe^3+^-siderophore outer membrane receptors revealed that FttA shows similarity to *E. coli fepA* (27% identity/44% similarity in amino acid level) and *Vibrio anguillarum fetA* (30% identity/48% similarity in amino acid level), suggesting its potential role as a Fe^3+^-turnerbactin uptake receptor. Although the TBDRss play an essential role for the iron uptake in bacteria, there are cases in which Fe^3+^-siderophores can be transported via multiple TBDRs encoded by genes that reside in different chromosomal loci (Poole et al., 1993;Rabsch et al., 1999;Mey et al., 2002;Ghysels et al., 2004;Hannauer et al., 2010;Naka and Crosa, 2012;Wyckoff et al., 2015;Payne et al., 2016). To investigate the role of the *fttA* gene, we constructed an in-frame *fttA* deletion mutant, and the growth of the *fttA* mutant was compared with the wild type strain and turnerbactin biosynthetic mutant (Δ*tnbF*), under iron rich and limiting conditions. As shown in **Figure 2,** the Δ*fttA* mutant did not grow under the iron limiting condition as compared with the wild type strain while this mutant still grew well in the iron rich growth condition. The growth of the Δ*fttA* mutant under the iron limiting condition was recovered when the *fttA* gene was expressed *in trans* in the *fttA* mutant confirming that the growth defect was due to the deletion of the *fttA* gene (**Figure 3**). These results indicate that the *fttA* gene is essential for the growth of *T. turnerae* under iron limiting conditions.

**Figure 1.**
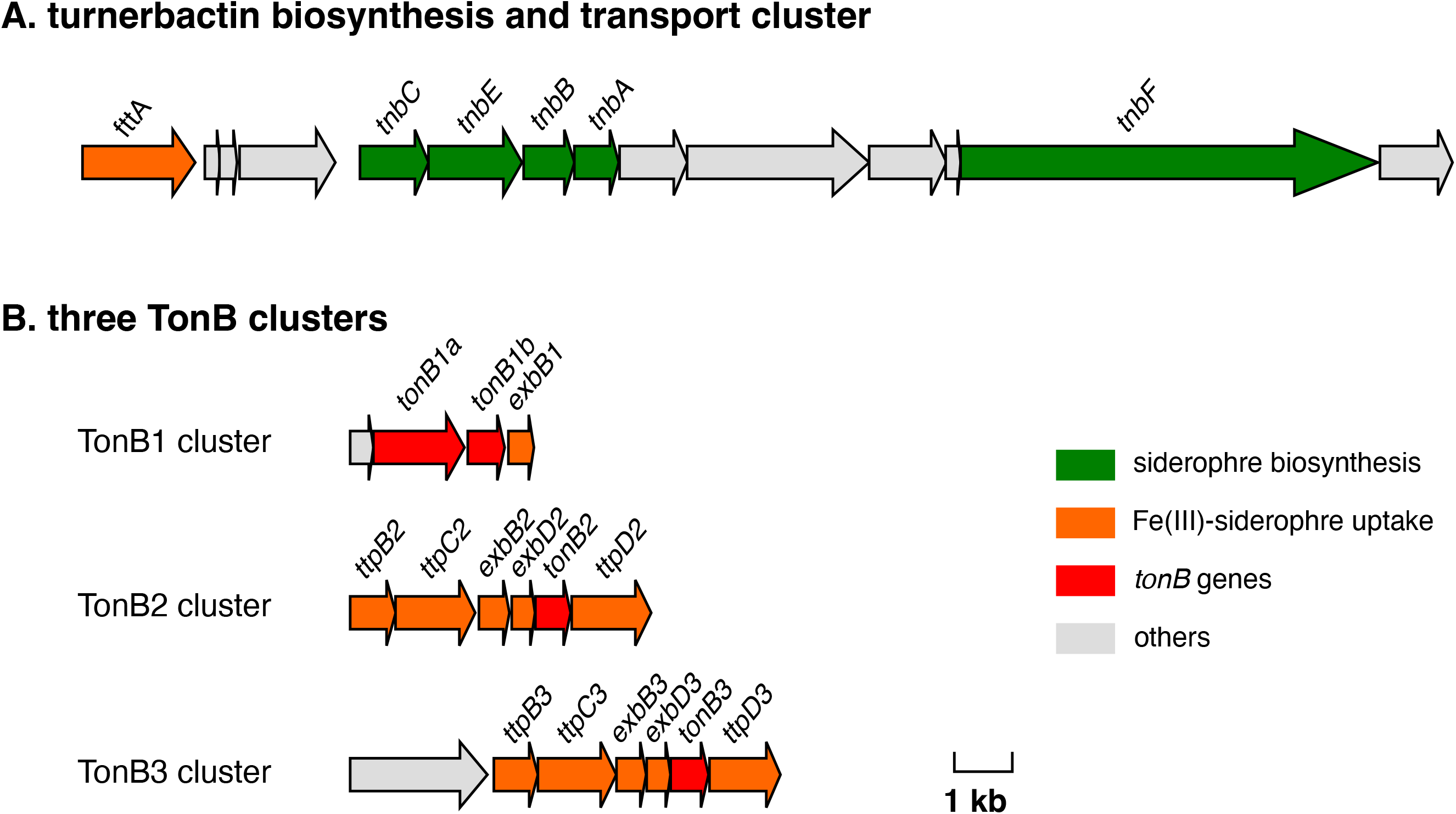
Gene clusters involved in siderophore-mediated iron transport. Panel A. the turnerbactin biosynthesis and transport cluster. Panel B. Three *tonB* clusters. The figure was modified from a gene cluster map constructed by Gene Graphics (Harrison et al., 2018). Green arrows indicate genes already characterized or predicted to be responsible for siderophore biosynthesis. Orange arrows indicate genes annotated to be involved in Fe(III)-siderophore uptake, and of those, TonB-homologues are shown in red color.

**Figure 2.**
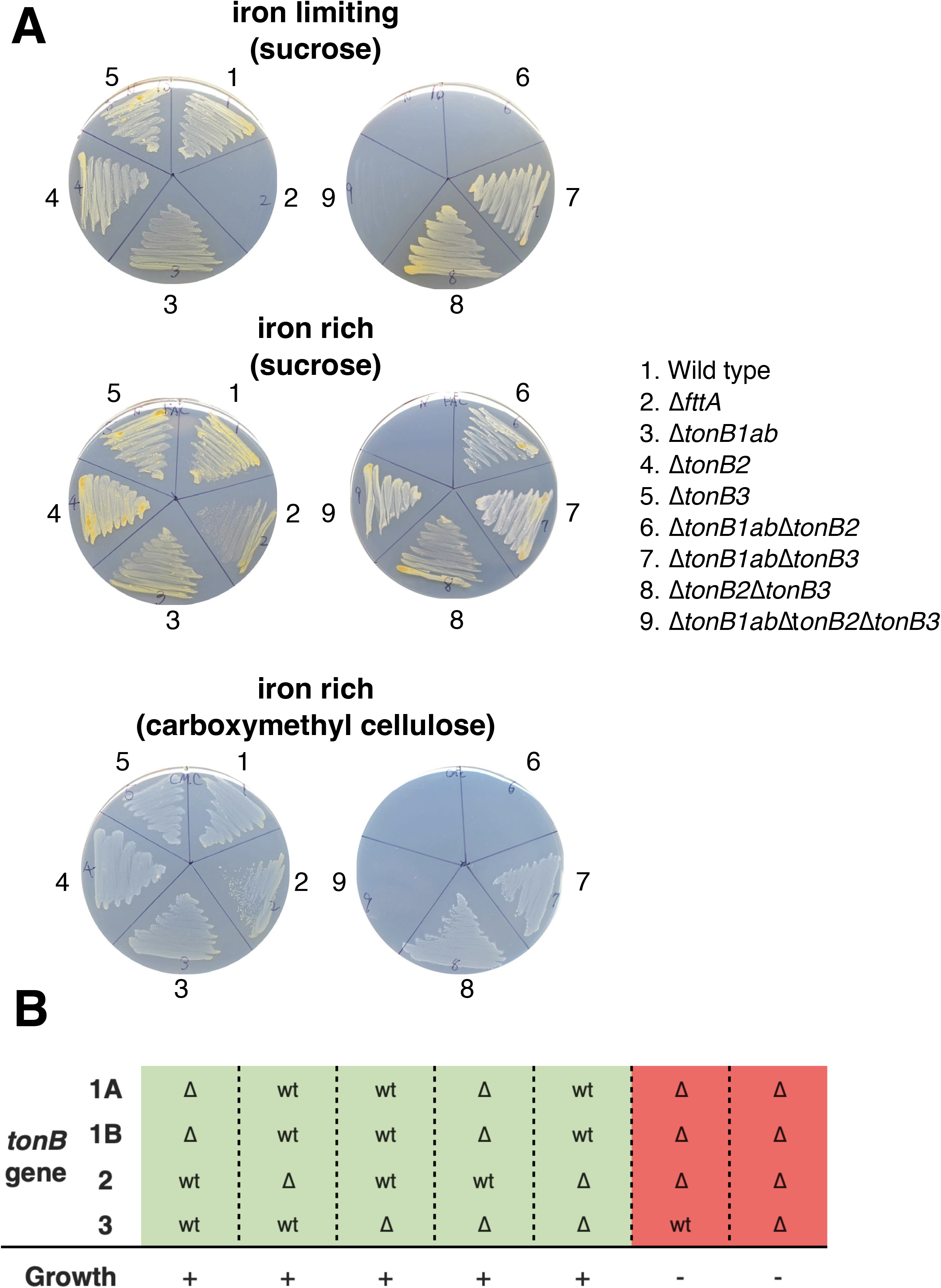
Involvement of *fttA* and TonB genes in iron transport and carbohydrate derived from cellulose. Panel A, Growth of *T. turnerae* mutants under different growth conditions. Sucrose (0.5 %) or carboxymethyl cellulose (0.5 %) were added as a sole carbon source in the SBM agar plates, and FAC (1μM) and EDDA (10 μM), and FAC (10 μM) were supplemented in the SBM medium to obtain iron limiting and rich conditions, respectively. *T. turnerae* strains were streaked on the plates, and the pictures were taken after 7 days incubation at 30 °C. FAC, ferric ammonium citrate; EDDA, ethylenediamine-N,N’-bis(2-hydroxyphenylacetic acid) Panel B, Growth response of *tonB* deletions to iron restriction and carboxymethyl cellulose combined. “wt” indicates the presence of wild type *tonB* genes while “Δ” shows the absence of the tonB gene (the in-frame gene deletion). The strains that grew under iron limiting conditions when sucrose was used as a carbon source or when carboxymethyl cellulose was used as a carbon source under iron rich conditions were highlighted as green and shown as “+” while the strains that did not grow were highlighted as red and shown as “−”;

**Figure 3.**
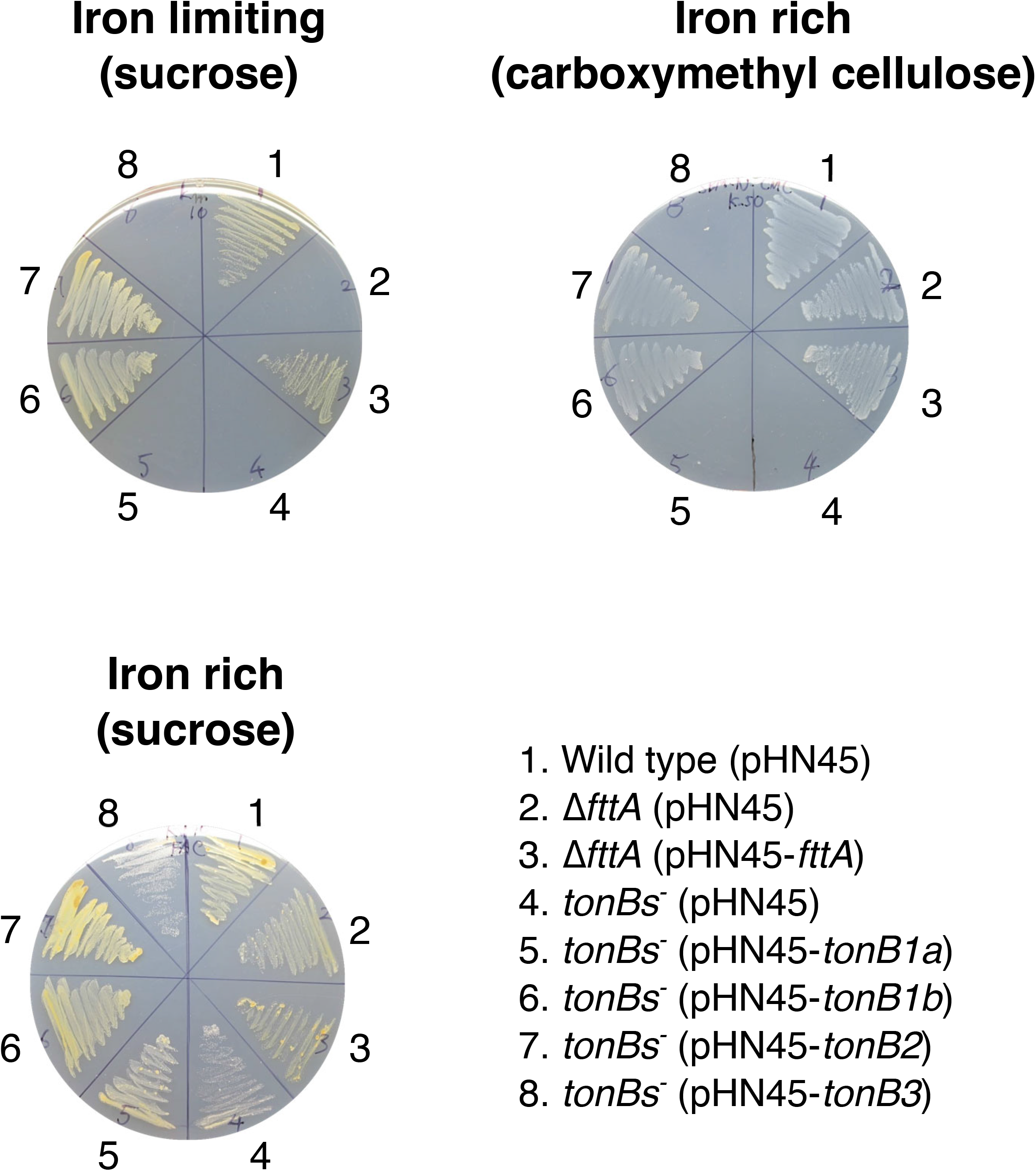
Complementation of *fttA* and *tonB* mutants. Sucrose or carboxymethyl cellulose were added as a sole carbon source in the SBM agar plates. FAC (1μM) and EDDA (10 μM), and FAC (10 μM) were added in the SBM medium to obtain iron limiting and rich conditions, respectively. *T. turnerae* strains were streaked on the plates, and the pictures were taken after 7 days incubation at 30 °C. FAC, ferric ammonium citrate; EDDA, ethylenediamine-N,N’-bis(2-hydroxyphenylacetic acid. pHN45, plasmid expression vector; tonBs-, Δ*tonB1ab*Δ*tonB2*Δ*tonB3*

To further investigate whether the growth deficiency of the *ΔfttA* mutant is due to the failure of Fe^3+^-turnerbactin uptake, we performed a bioassay (siderophore cross-feeding assay). We first constructed a turnerbactin production deficient strain, Δ*tnbA*Δ*tnbF*. The *tnbA* gene was also mutated to eliminate the 2,3-dihydroxybenzoate-2,3-dehydrogenase (2,3-DHBA) production since 2,3-DHBA also acts as an iron chelator (Bellaire et al., 2003). The *fttA* gene was mutated in the Δ*tnbA*Δ*tnbF* background. Supplementation of the iron chelator, ethylenediamine-di-(o-hydroxyphenyl acetic acid) (EDDA) into growth medium led to the failure of the growth of the turnerbactin production deficient strains, Δ*tnbA*Δ*tnbF* and Δ*tnbA*Δ*tnbF*Δ*fttA* (**Figure 4**). This growth defect of Δ*tnbA*Δ*tnbF* was overcome when the wild type strain producing turnerbactin was spotted on the agar plate containing Δ*tnbA*Δ*tnbF* (see the growth halo around the spot). However, the Δ*tnbA*Δ*tnbF*Δ*fttA* strain in which the *fttA* gene was deleted didn’t recover its growth in the presence of the wild type strain spot while spotting ferric ammonium citrate was able to recover its growth. These results indicate that the *fttA* gene is essential for the uptake of turnerbactin produced by the wild type strain. Furthermore, to test whether *T. turnerae* T7901 can utilize an exogenous siderophore produced by marine bacterium *Vibrio campbellii*, we used extracts obtained from wild type *V. campbellii* that produces amphi-enterobactin and anguibactin, and its derivatives, an amphi-enterobactin producer and an anguibactin producer (Naka et al., 2018). Extracts rather than cultures were used because *V. campbellii* strains cannot grow on SBM medium. The growth of *T. turnerae* was recovered when amphi-enterobactin was provided by the indicator strain while anguibactin was not able to compensate for the growth defect under iron limiting conditions. These results indicate that *T. turnerae* can take up amphi-enterobactin but not anguibactin produced by the marine pathogenic bacterium *V. campbellii*.

**Figure 4.**
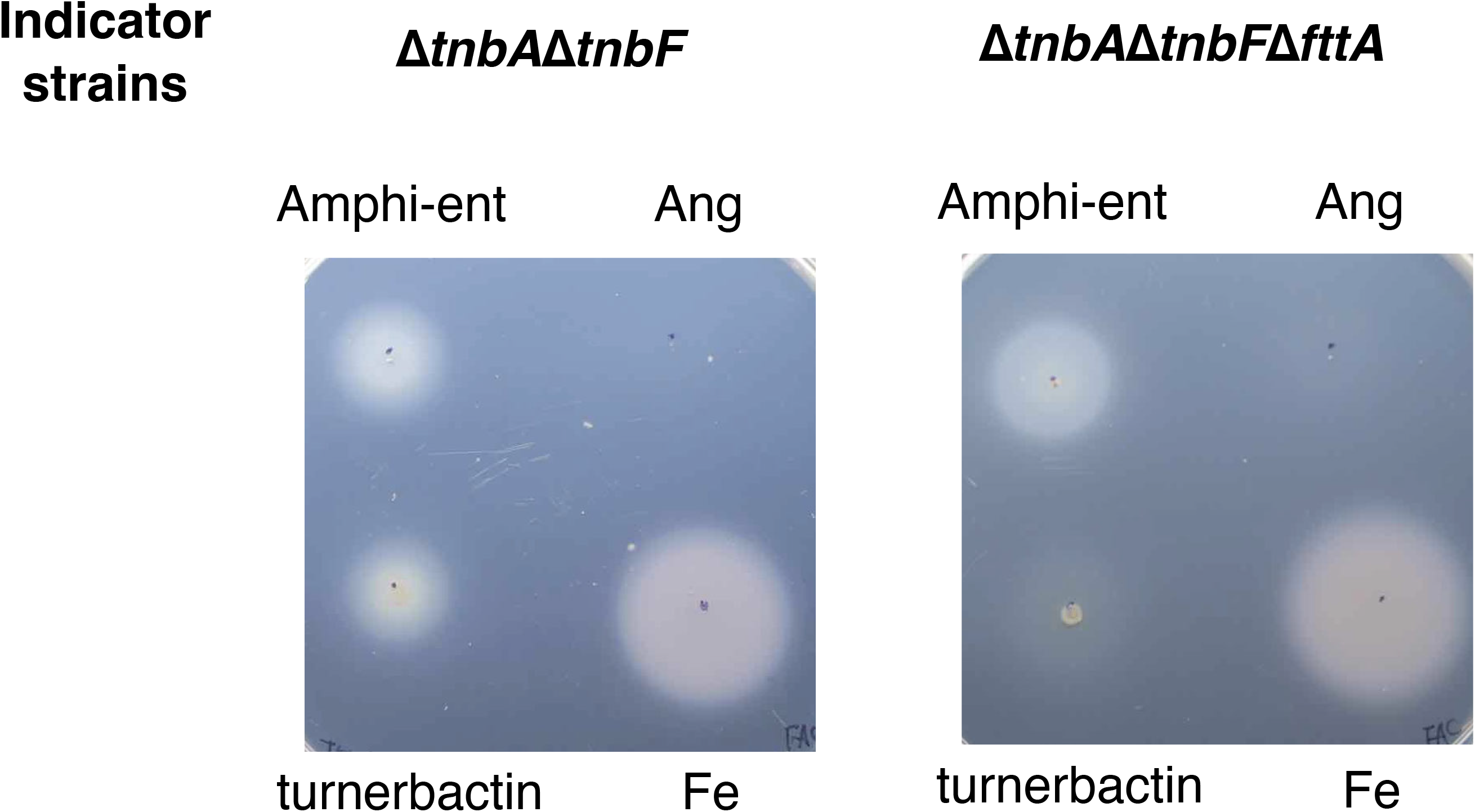
Bioassay to test the involvement of FttA in transport of endogenous and exogenous siderophores. Δ*tnbA*Δ*tnbF* produces neither turnerbactin nor its precursor due to the mutation in *tnbF* and *tnbA*, respectively, therefore cannot grow under iron limiting growth condition generated by supplementing an iron chelator, ethylenediamine-N,N’-bis(2-hydroxyphenylacetic acid. *T. turnerae* strains were grown under non-aggregation conditions. T7901, 5 μl of *T. turnerae* T7901 culture (containing turnerbactin); Amphi-ent, extracts containing amphi-enterobactin obtained from *V. campbellii* HY01Δ*angR* (Naka et al., 2018); extracts contain anguibactin prepared from *V. campbellii* HY01Δ*aebF* (Naka et al., 2018); Fe, 5μl of ferric ammonium citrate. Pictures were taken after 7 days incubation at 30°C.

### Identification of TonB clusters in *T. turnerae* T7901

The presence of the TonB2 and TonB3 clusters in *T. turnerae* was briefly described before and those are similar to the TonB2 and TonB3 clusters of marine vibrios such as *V. vulnificus* (Kuehl and Crosa, 2010), but the function of those *tonB* genes has not been elucidated yet. By sequence similarity searching of protein sequences annotated in *T. turnerae* T7901 with well-characterized TonB genes from *E. coli* K-12, and marine bacteria including *V. vulnificus, V. cholerae, V. anguillarum* and *Aeromonas hydrophila*, we identified two more *tonB* gene homologues in a cluster (named here TonB1 cluster) in addition to TonB2 and TonB3 clusters. Interestingly, the TonB1 cluster carries two TonB genes, *tonB1a* and *tonB1b*, located next to each other and an *exbD* gene homologue (*exbD1*), but an *exbB* homologue was not found in this cluster (**Figure 1**). The TonB1 clusters in vibrios (consisting of *tonB1, exbB1* and *exbD1*) are located linked to the heme/hemoglobin transport cluster (Occhino et al., 1998;O’Malley et al., 1999;Stork et al., 2004;Wang et al., 2008;Kustusch et al., 2012). However, there is no heme cluster near the TonB1 system in *T. turnerae*. Prediction of transmembrane helices with TMHMM server version 2 (https://services.healthtech.dtu.dk/service.php?TMHMM-2.0) (Krogh et al., 2001) indicated that TonB1b, TonB2 and TonB3 harbor one transmembrane domain typically found in classical TonB proteins while TonB1a is an unusual TonB protein that carries an extended N-terminal domain predicted to carry four transmembrane domains, that can be found small number of bacteria (Chu et al., 2007).

### Two TonB genes are essential for the growth of *T. turnerae* under iron limiting growth conditions

To understand which TonB gene(s) facilitate the growth of *T. turnerae* under specified conditions, single and multiple *tonB* gene mutants were constructed. Since *tonB1a* and *tonB1b* genes are co-located, both *tonB1a* and *tonB1b* were deleted together, generating the *ΔtonB1ab* mutant. The growth of those strains was compared under iron rich and limiting growth conditions. As shown in **Figure 2**, single mutants that lack *tonB* gene(s) in each TonB cluster such as *ΔtonB1ab, ΔtonB2* and *ΔtonB3* as well as the double *tonB* gene mutants in the TonB1 and TonB3 cluster (*ΔtonB1abΔtonB3*) and in the TonB2 and TonB3 cluster (*tonB2ΔtonB3*) showed growth under both iron rich and limiting growth conditions. On the other hand, the *tonB* gene mutants in both the TonB1 and TonB2 cluster, *ΔtonB1abΔtonB2* and the quadruple *tonB* gene mutant in which all *tonB* genes were eliminated, *ΔtonB1abΔtonB2ΔtonB3*, did not grow under iron limiting conditions. Similar results were observed in the turnerbactin biosynthetic deficient mutant *ΔtnbF* and the ferric-turnerbactin transport deficient *ΔfttA* mutant. These results indicate that the *tonB* genes in both the TonB1 and TonB2 cluster are involved in the iron transport in *T. turnerae* T7901.

We further performed complementation experiments to confirm that the growth defect of some of mutants were not due to polar effects and/or secondary mutations, and also to understand which *tonB1* genes (*tonB1a* or *tonB1b*) is responsible for the growth of *T. turnerae* T7901 under iron limiting conditions. *tonB* genes with their ribosomal binding sites were cloned in the expression vector pHN33, and conjugated into the *ΔtonB1abΔtonB2ΔtonB3* mutant. The expression of all four TonB genes was confirmed by RT-PCR (**Figure S1**). As shown in **Figure 3**, the growth of the quadruple *tonB* mutant under iron limiting growth conditions were recovered only when *tonB1b* or *tonB2* genes are expressed *in trans* in the quadruple *tonB* mutant. All strains grew well under an iron rich growth condition. From these results, we conclude that out of four *tonB* genes, *tonB1b* and *tonB2* are responsible for the growth of *T. turnerae* T7901 under iron limiting conditions, and *tonB1a* and *tonB3* are not responsible for iron uptake under this growth condition.

### Involvement of TonB genes in the growth of *T. turnerae* T7901 cellulose as a carbon source

During the course of mutant construction in TonB genes, it was very hard to obtain the *ΔtonB1abΔtonB2* mutant. We realized that this *ΔtonB1abΔtonB2* mutant does not grow when cellulose is used as a sole carbon source in the growth medium. This mutant did not show a growth defect on sucrose plates. Since supplementation of cellulose and carboxymethyl cellulose (cellulose derivative) into the growth medium resulted in same consequences, we decided to use carboxymethyl cellulose due to its solubility in growth medium. We further tested the growth of all single and multiple *tonB* gene mutants on SBM medium supplemented with either sucrose or carboxymethyl cellulose as a carbon source, and we found that the mutants missing *tonB* genes in the both TonB1 and TonB2 clusters (*ΔtonB1abΔtonB2*) and the strain that lacks all *tonB* genes (*ΔtonB1abΔtonB2ΔtonB3*) showed a dramatic growth defect when carboxymethyl cellulose was used as a sole carbon source (**Figure 2**). The rest of the mutants tested grew on both sucrose and cellulose media. The growth defect in the quadruple TonB mutant, *ΔtonB1abΔtonB2ΔtonB3* was recovered when the *tonB1b* or *tonB2* genes were expressed *in trans* in the mutant while *tonB1a* and *tonB3* was not able to compensate the growth defect on cellulose plates (**Figure 3**). These results demonstrate that *tonB1b* and *tonB2* are involved in carbohydrate utilization when cellulose is provided as a sole carbon source. Turnerbactin biosynthesis (*ΔtnbF*) and transport (*ΔfttA*) deficient mutants did not show growth defects on cellulose plates, therefore the growth defect appears to be independent of turnerbactin production and utilization.

### Iron regulation of turnerbactin biosynthesis and transport genes

It has been proposed that Region 7 consists of two iron-regulated transcriptional units and both operons might be regulated by the ferric uptake regulator since two possible Fur binding sites (Fur boxes) were identified in the upstream regions of *fttA* and *tnbC* (Han et al., 2013). We performed RT-qPCR analysis to test whether genes in Region 7 are actually iron-regulated. Our results clearly showed that three representative genes such as *tnbA, tnbF* and *fttA* are up-regulated under iron limiting growth conditions (**Figure 5**). We also performed a Fur titration assay to test whether the *E. coli* ferric uptake regulator (Fur) binds to these putative TonB boxes. The result in **Figure S2** shows that *E. coli* Fur can bind to two Fur boxes as compared with two negative controls, indicating that Fur is involved in the up-regulation of those genes under iron limiting conditions.

**Figure 5.**
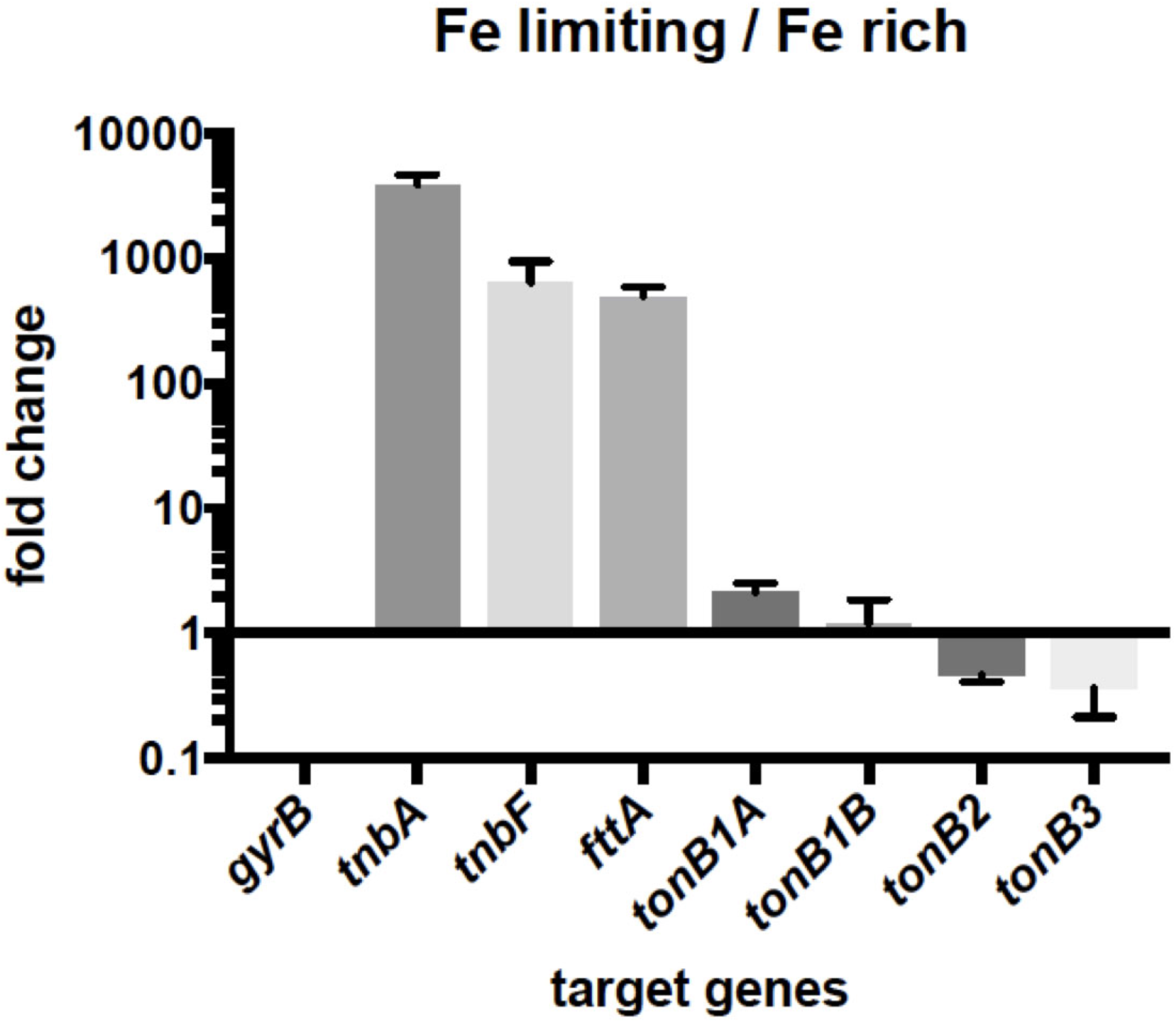
Regulation of iron transport-related genes in *T. turnerae* T7901. Expression of genes between iron limiting and rich growth conditions were compared by qRT-PCR. The data represents the mean value of at least three biological replicates with error bars that are the standard error of the mean.

### Iron regulation of TonB genes

In many bacteria, TonB genes are typically regulated by iron to control internal iron concentration (Young and Postle, 1994;Zhao and Poole, 2000;Bosch et al., 2002;Ochsner et al., 2002;Bjarnason et al., 2003;Beddek et al., 2004;Osorio et al., 2004;Stork et al., 2004;Mey et al., 2005;Bender et al., 2007;Wang et al., 2008). To test iron-regulation of TonB genes in *T. turnerae*, we performed RT-qPCR using primers to amplify each TonB gene (**Figure 5**). The results indicate that relative expression levels of all TonB genes were not changed as much as those of *tnbA* and *tnbF* which were dramatically increased under iron limiting conditions as compared with iron rich conditions.

## Discussion

One of the compounds *Teredinibacter turnerae* produces is the siderophore turnerbactin that is used to acquire iron which is an essential metal for their growth in iron limiting environments. It has been suggested that turnerbactin might be used to compete for iron with casually associated environmental bacteria to survive under iron limiting conditions which are typically found in marine environments and inside hosts (Han et al., 2013). Turnerbactin-related genes were found in the secondary metabolite cluster, Region 7 located within GCF_8 (identified by metagenomics), and the *tnbF* gene is essential for turnerbactin biosynthesis (Han et al., 2013;Altamia et al., 2020). However, the transport mechanism of Fe(III)-turnerbactin was not characterized yet. To transport Fe(III)-siderophore across the outer membrane, Gram negative bacteria require the TonB system that typically consists of TonB, ExbB and ExbD, that transduce proton motive force generated in the inner membrane to outer membrane receptors, resulting in conformational change in the outer membrane receptors (Ratliff et al., 2022). The TonB system was originally found and has been extensively characterized in *E. coli* (Postle and Larsen, 2007). *E. coli* and many other bacteria carry a single set of the TonB system, but after finding two TonB systems in *V. cholerae* (Occhino et al., 1998), multiple TonB systems have been identified and characterized in a number of bacteria, including many *Vibrio* species (2 or 3 systems) (Seliger et al., 2001;Stork et al., 2004;Alice et al., 2008;Wang et al., 2008;Tanabe et al., 2012), *Aeromonas hydrophila* (3 systems) (Dong et al., 2016;Dong et al., 2019;Dong et al., 2023), *Pseudomonas aeruginosa* (3 systems) (Poole et al., 1996;Zhao and Poole, 2000;Huang et al., 2004), *Acinetobacter baumannii* (3 systems) (Zimbler et al., 2013;Runci et al., 2019), and *Bacteroides fragilis* (6 systems) (Parker et al., 2022). In those examples, some TonB systems are functionally independent while others show functionally redundancy, for transport for particular substances such as siderophores and other nutrients, or physiological activities.

The aim of this study is to explain the Fe(III)-turnerbactin uptake mechanism. The *fttA* gene located in Region 7 is a homologue of Fe(III)-siderophore TonB-dependent outer membrane receptors (TBDRs). RT-qPCR analysis showed that the *fttA* gene as well as two turnerbactin biosynthetic genes, *tnbA* and *tnbF* are clearly up-regulated under iron limiting growth conditions. Iron-regulation of genes in Region 7 were further analyzed by RNA sequencing (RNA-seq), and all annotated iron transport-related genes in Region 7, *fttA* to *TERTU_RS18085* are up-regulated under iron limiting conditions (**Table S3**). Furthermore, the Fur titration assay (FURTA) showed that the *E. coli* ferric uptake regulator, Fur, can bind to the potential promoter regions previously identified and located upstream of *fttA* and *tnbC* whereas the upstream region of *TERTU_RS18075* showed a negative result. Taken together, the iron regulation of Region 7 is caused by at least two distinct promoters in a Fur-dependent manner, as proposed before (Han et al., 2013). We constructed an in-frame deletion mutant of *fttA*, and showed that the *fttA* gene is responsible for Fe(III)-turnerbactin transport and indispensable for growth under iron-limiting conditions while the *fttA* mutant grew well under iron-rich growth conditions, demonstrating that FttA is the sole TBDR involved in Fe(III)-turnerbactin uptake. We also tested the ability of *T. turnerae* to transport xenosiderophores, amphi-enterobactin and anguibactin, from *Vibrio campbellii*. Our results showed that *T. turnerae* can utilize Fe(III)-amphi-enterobactin or its hydrolyzed derivatives as an iron source and it was independent of *fttA*, whereas Fe(III)-anguibactin failed to enhance the growth of *T. turnerae* under iron limiting conditions. Amphi-enterobactin is produced by both *V. campbellii* and *V. harveyi* that are members of the Harveyi clade ubiquitously found in marine environments and some strains are causative agents of vibriosis that affect marine vertebrates and invertebrates. On the other hand, anguibactin is produced by *V. campbellii* but not by *V. harveyi* (Zane et al., 2014;Naka et al., 2018). Our findings indicate that *T. turnerae* possess the ability to “steal” iron from the siderophore or its derivatives commonly found in different species rather than from the species specific siderophore, and this might provide an advantage to *T. turnerae* to survive in marine environments where amphi-enterobactin is available. It is still unknown what gene(s) is encoding the outer membrane receptor for Fe(III)-amphi-enterobactin since *fttA* was not required for Fe(III)-amphi-enterobactin utilization. In the *T. turnerae* T7901 genome, 38 genes were annotated to encode TonB-dependent outer membrane receptors, and our RNA-seq result indicated that 6 genes in addition to *fttA* were up-regulated (logFC > 1) under iron-limiting growth conditions (**Table S4**), and one or some of them might be responsible for Fe(III)-amphi-enterobactin utilization.

By searching in the genome of *T. turnerae*, we identified four TonB genes that are located in three TonB clusters. The *tonB1* cluster of *T. turnerae* is a unique *tonB* cluster that contains two *tonB* genes, *tonB1a* and *tonB1b*, and *exbD1*, but lacks *exbB* typically found in TonB clusters. TonB1a carries a N-terminal extension as compared with conventional TonB proteins, and this type of TonB protein was identified by bioinformatic analysis but the function is still unknown (Chu et al., 2007). The gene organization of TonB2 and TonB3 clusters resemble marine vibrios and contain homologues of *ttpB, ttpC, exbB, exbD, tonB* and *ttpD* in which *ttpB, ttpC* and *ttpD* are specifically found in vibrios and some marine bacteria (Kuehl and Crosa, 2010;Barnes et al., 2020). In vibrios, the TonB2 cluster is involved in iron transport while the function of the TonB3 cluster is still unknown (Alice and Crosa, 2012;Duong-Nu et al., 2016). The similarity of TonB2 and TonB3 systems, especially the presence of *ttpB, ttpC* and *ttpD*, to those of vibrios indicate that the *tonB2* system could provide benefits to adapt in coastal waters where both *T. turnerae* and vibrios live. Conversely, the TonB1 cluster of *T. turnerae* did not show similarity to that of vibrios. *Vibrio* TonB1 systems are linked to gene clusters that are responsible for hemin/hemoglobin uptake, and are involved in hemin/hemoglobin uptake (Occhino et al., 1998;O’Malley et al., 1999;Mourino et al., 2004;Lemos and Osorio, 2007;Wang et al., 2008;Septer et al., 2011;Kustusch et al., 2012). We speculate that *T. tunerae* did not evolve a similar TonB1 cluster possibly due to the absence of a heme/hemoglobin cluster, and *T. turnerae* does not encounter environments in which hemin and/or hemoglobin is available during their life cycle, due to the absence of hemoglobin in bivalves such as the shipworm hosts.

In most bacteria, TonB genes are normally up-regulated in iron limiting conditions (actually repressed under iron rich conditions) because excess amounts of iron are toxic to bacteria because it leads to Fenton reaction causing the overproduction of reactive oxygen species in presence of oxygen. Interestingly, RT-qPCR results showed that none of TonB genes as well as other genes in *tonB* clusters are clearly regulated under iron limiting conditions and the expression pattern of TonB genes were further confirmed by RNA-seq, supporting the result of RT-qPCR and also suggesting that the regulation occurs at a cluster level. It has been reported that *tonB3* genes in *V. vulnificus* and *A. hydrophila* are not iron regulated (Alice and Crosa, 2012;Dong et al., 2019). However, neither of the *tonB3* genes in those bacteria are involved in iron transport. It is of interest that all *T. turnerae* TonB genes are not clearly iron-regulated even though *tonB1b* and *tonB2* genes *are* involved in Fe(III)-turnerbactin utilization. These results indicated that *T. turnerae* might still need *tonB* genes expressed even in iron rich conditions.

One of the unusual features of *T. turnerae* is its ability to degrade lignocellulose from wood and utilize its derivatives as a carbon source (Waterbury et al., 1983;Distel et al., 2002b). It has been reported that some bacteria use TBDRs to take up plant-derived carbohydrates, mono- and poly-saccharides. *Xanthomonas campestris* pv. *campestris* (Xcc) use TBDR to take up sucrose, and the comparative genomic and gene expression analysis suggested that Xcc as well as some marine bacteria possibly take up plant carbohydrates via TBDRs (Blanvillain et al., 2007). *Caulobacter crescentus* uses the TonB1 system to transport maltose and maltodextrins (Neugebauer et al., 2005;Lohmiller et al., 2008). We showed that two of the *tonB* genes, *tonB1b* and *tonB2*, are involved in carbohydrate utilization derived from cellulose in *T. turnerae* whereas mutations in other *tonB* genes (*tonB1a* and *tonB3*) did not affect the growth. These results indicate that the same set of *tonB1b* and *tonB2* are functional not only for Fe(III)-turnerbactin uptake but also cellulose utilization. Further studies are required to identify TBDRs involved in the uptake of cellulose-derived carbohydrates, and what carbohydrate(s) are transported across the outer membrane. Some TBDRs are located close to genes potentially involved in hemicellulose degradation (Data not shown). It is worth noting that the experiments were performed under iron rich conditions, therefore the lack of growth was not due to iron limitation. These results indicate that *tonB1b* and *tonB2* are functional even under iron rich conditions. All *tonB* genes are expressed in both iron rich and limiting growth conditions (**Table S3**). This dual role of TonB1b and TonB2 for iron and carbohydrate uptake could explain why *T. turnerae* does not clearly regulate those genes depending on iron concentrations although TonB genes are down-regulated under iron rich conditions in most bacteria.

In summary, the *fttA* gene, a homologue of Fe(III)-siderophore TBDR genes, is indispensable for the survival of *T. turnerae* under iron limiting growth conditions because it is essential for Fe(III)-turnerbactin utilization as an iron source. FttA appears to be essential for the transport of Fe(III)-turnerbactin across the outer membrane, and Fe(III)-amphi-enterobactin produced by other marine bacteria can be utilized as an iron source without FttA. Two out of four *tonB* genes, *tonB1b* and *tonB2*, show functional redundancy for the survival of *T. turnerae* under iron limiting conditions as well as the growth of *T. turnerae* when cellulose was supplied as a sole carbon source. Since *tonB* genes are known to energize TBDRs to substrate import across the outer membrane, those findings indicate that carbohydrates derived from cellulose are likely transported by TBDRs. All of genes in Region 7 encompassing *fttA* to *TERTU_RS18085* were repressed under iron rich conditions to avoid intracellular excess iron whereas the expression of the *tonB* genes is remained under iron rich conditions, indicating the importance of *tonB* genes even under iron rich conditions possibly for the utilization of cellulose as a carbon source.

## Acknowledgements

This work was supported by National Institutes of Health Grant 1U01TW008163. We would like to thank Drs. Aaron Puri (University of Utah), Daniel Distel (Northeastern University) and Alison Butler (University of California, Santa Barbara) for valuable comments.

---

**Table S1.**
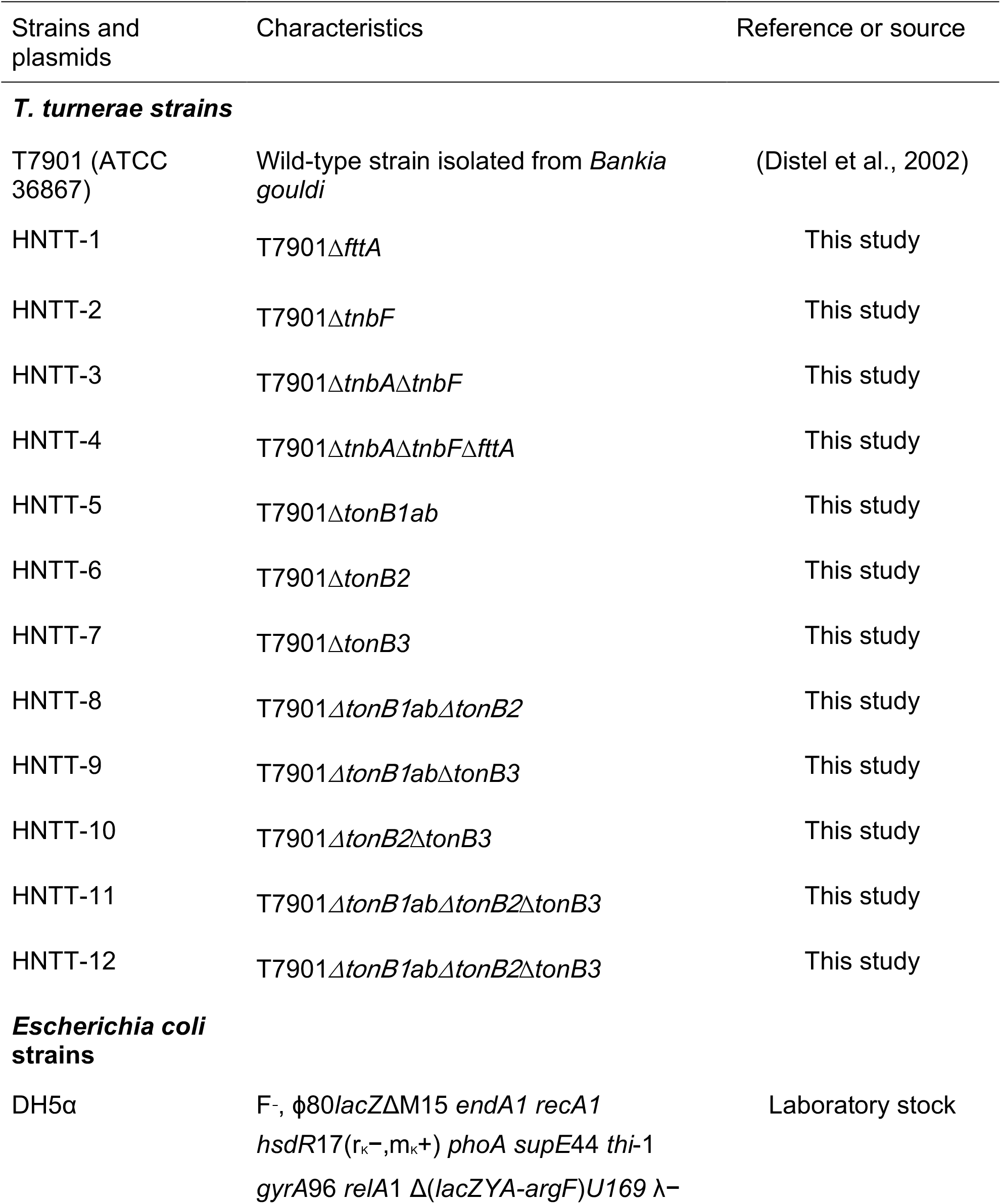
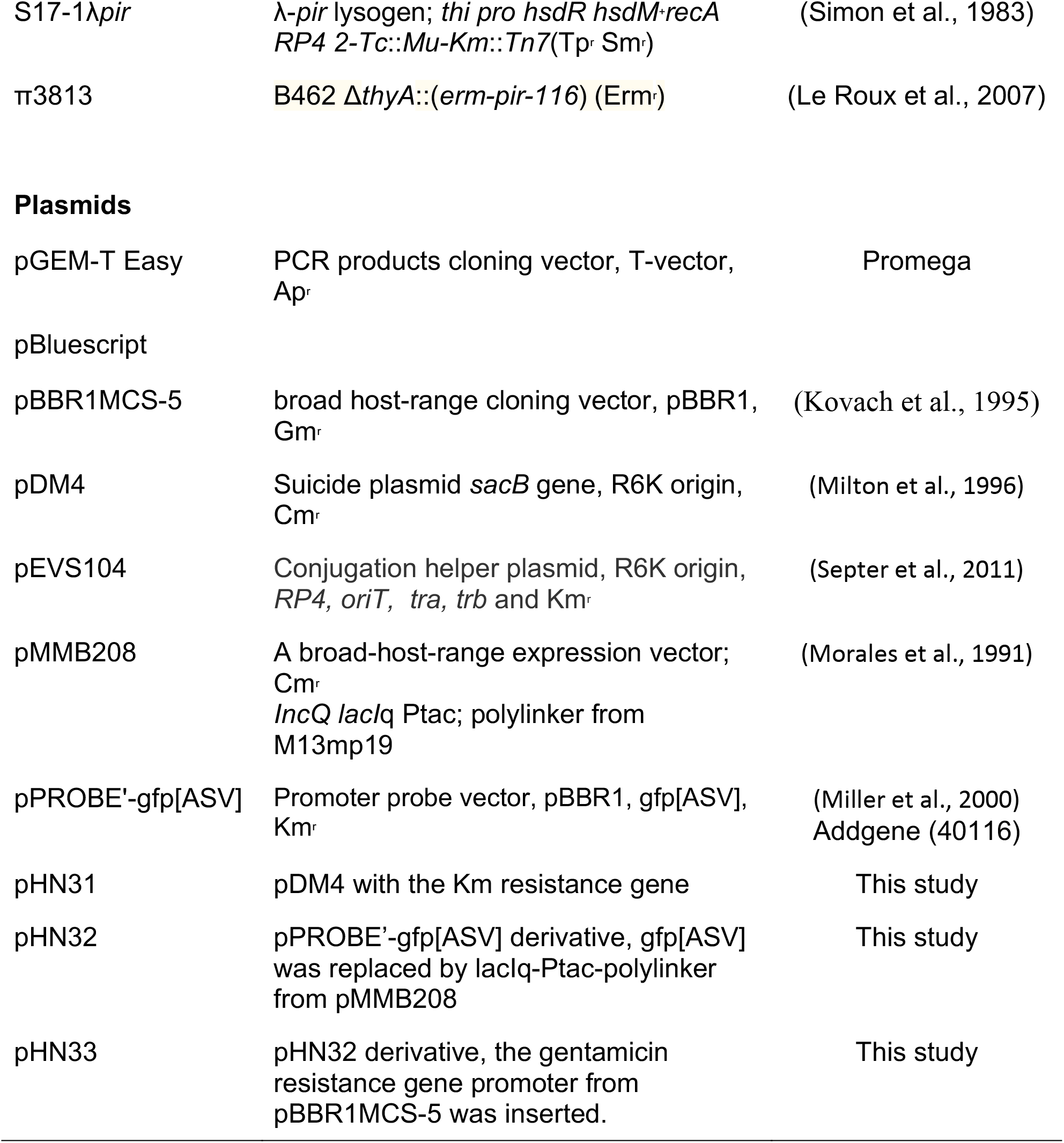
Strains and plasmids used in this study

**Table S2.**
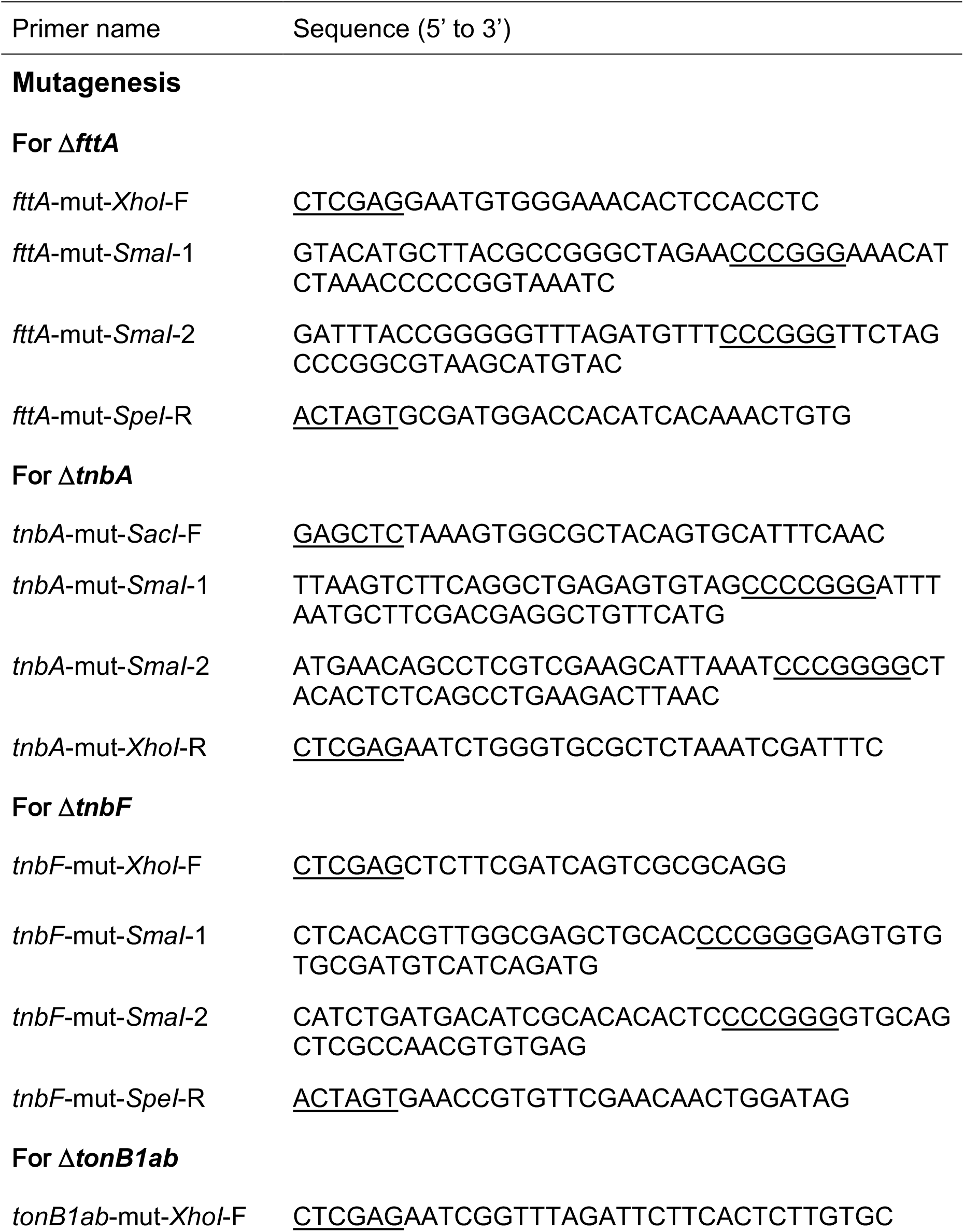
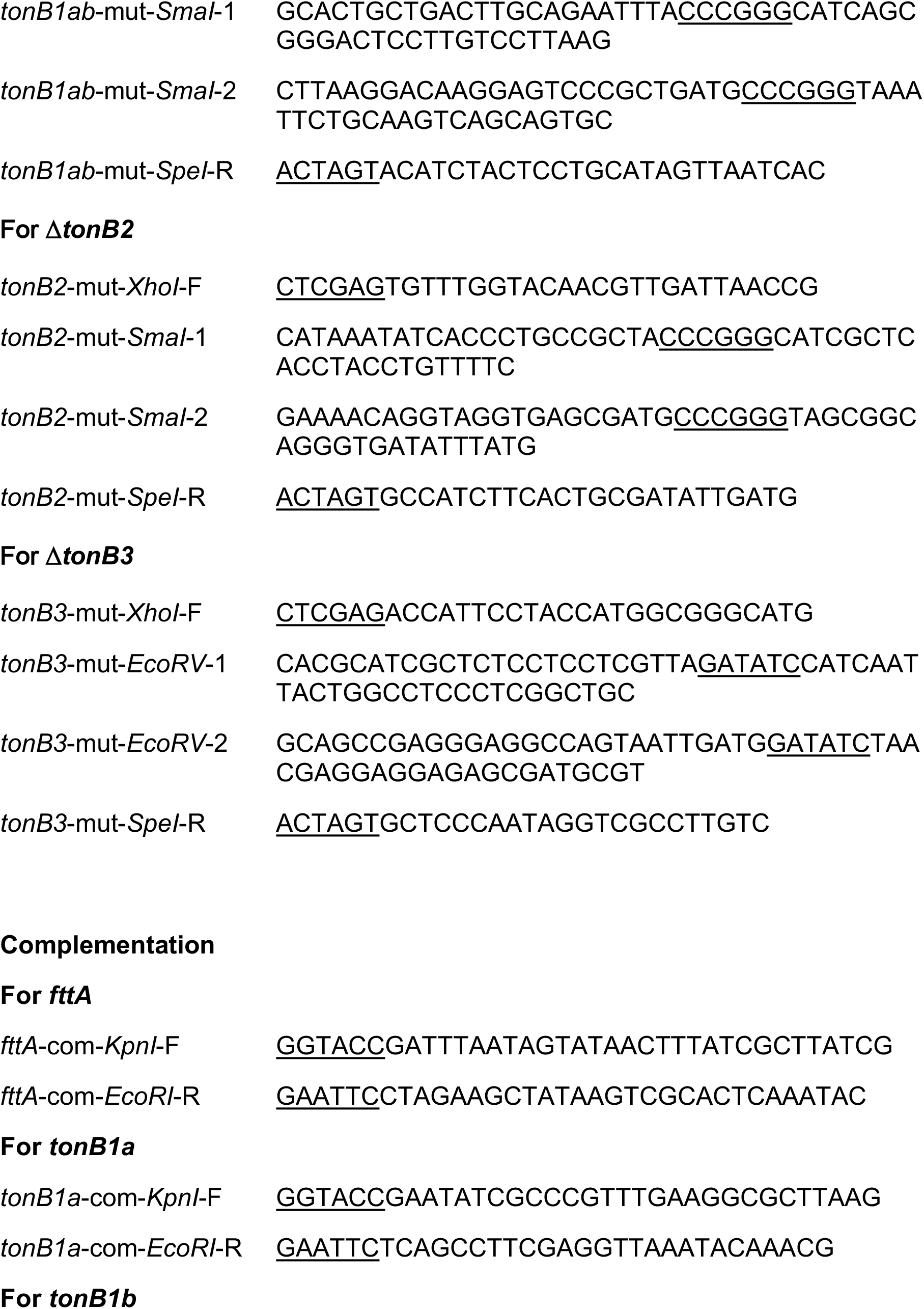
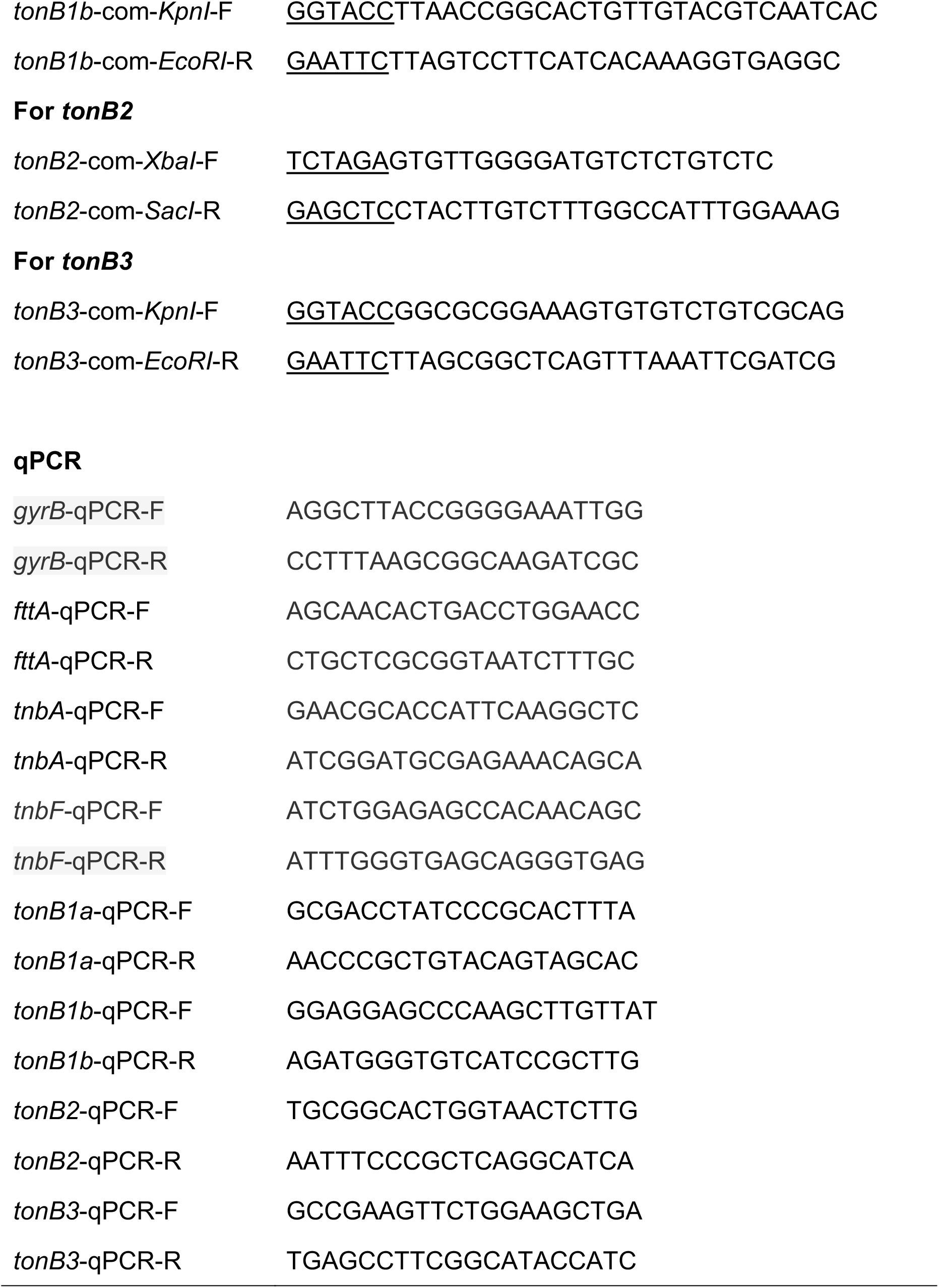
Primers used in this study

## RNA-sequencing

*T. turnerae* T7901 was grown in non-aggregation SBM broth medium containing sucrose supplemented with 10 μM FAC (iron rich condition) or 0.1 μM FAC (iron limiting condition), and the expression of genes were compared by RNA-sequencing (RNA-seq). We avoided using an iron chelator to exclude any effects other than iron chelation caused by iron chelators. 2 biological replicates were used.

Illumina library construction using the Illumina TruSeq Stranded Total RNA Library Prep Kit with Ribo-Zero (Illumina) and sequencing using Novaseq 6000 with 150 bp paired-end runs was performed at the Huntsman Cancer Institute’s High-Throughput Genomics Center at the University of Utah. All raw sequencing reads were deposited in the NCBI Sequence Read Archive (SRA) under the accession number PRJNA885807.

Quality control and preprocessing of FASTQ reads were performed using the fastp program (Chen et al., 2018). The processed reads were mapped on the chromosome of *T. turnerae* T7901 using HISAT2 (Kim et al., 2019) and the output was converted to bam files using SAMtools (Li et al., 2009). FeatureCounts (Liao et al., 2014) were used to count the reads on *T. turnerae* T7901 genes, and differential expression analysis was carried out using the EdgeR package (Robinson et al., 2010).

---

**Table S3.** RNA-seq analysis to understand iron regulation of iron transport genes Turnerbactin biosynthesis and transport cluster

| **Turnerbactin biosynthesis and transport cluster** |  |  |  |  |
| --- | --- | --- | --- | --- |
| Locus tag | gene name or annotation | logFC | logCPM | FDR-adjusted p-value |
| TERTU_RS18025 | *fttA* | 8.55 | 12.09 | 2.94E-64 |
| TERTU_RS18030 | hypothetical protein | 7.13 | 7.10 | 1.42E-66 |
| TERTU_RS18035 | hypothetical protein | 7.39 | 7.23 | 9.20E-66 |
| TERTU_RS18040 | PepSY domain-containing protein | 6.19 | 9.74 | 9.50E-46 |
| TERTU_RS18045 | *tnbC* | 11.52 | 12.90 | 5.89E-93 |
| TERTU_RS18050 | *tnbE* | 12.27 | 13.20 | 5.32E-98 |
| TERTU_RS18055 | *tnbB* | 12.72 | 12.92 | 4.30E-98 |
| TERTU_RS18060 | *tnbA* | 11.24 | 11.69 | 2.03E-88 |
| TERTU_RS18065 | efflux RND transporter periplasmic adaptor | 10.83 | 11.60 | 1.20E-72 |
| TERTU_RS18070 | multidrug efflux RND transporter permease | 6.9 | 12.44 | 1.84E-44 |
| TERTU_RS18075 | esterase | 8.41 | 10.40 | 5.04E-65 |
| TERTU_RS18080 | MbtH domain protein | 9.61 | 9.60 | 5.90E-76 |
| TERTU_RS18085 | *tnbF* | 8.41 | 13.36 | 3.25E-60 |
| TERTU_RS18090 | *tnbS* | 8.56 | 11.54 | 1.23E-62 |

**TonB1 cluster**
| Locus tag | gene name or annotation | logFC | logCPM | FDR-adjusted p-value |
| --- | --- | --- | --- | --- |
| TERTU_RS04350 | transcriptional regulator | 0.31 | 2.96 | 0.44 |
| TERTU_RS04355 | *tonB1a* | 1.17 | 5.86 | 2.55E-03 |
| TERTU_RS04360 | *tonB1b* | 0.33 | 3.93 | 0.49 |
| TERTU_RS04365 | *exbB1* | -0.59 | 3.12 | 0.1 |

**TonB2 cluster**
| Locus tag | gene name or annotation | logFC | logCPM | FDR-adjusted p-value |
| --- | --- | --- | --- | --- |
| TERTU_RS01630 | *ttpB2* | -1.03 | 11.08 | 0.01 |
| TERTU_RS01635 | *ttpC2* | -0.81 | 11.59 | 0.05 |
| TERTU_RS01640 | *exbB2* | -0.74 | 10.02 | 0.09 |
| TERTU_RS01645 | *exbD2* | -0.76 | 9.50 | 0.09 |
| TERTU_RS01650 | *tonB2* | -0.51 | 9.78 | 0.26 |
| TERTU_RS01655 | *ttpD2* | -0.5 | 10.93 | 0.28 |

**TonB3 cluster**
| Locus tag | gene name or annotation | logFC | logCPM | FDR-adjusted p-value |
| --- | --- | --- | --- | --- |
| TERTU_RS08980 | TonB-dependent receptor | -0.42 | 4.39 | 0.21 |
| TERTU_RS08985 | *ttpB3* | -1.61 | 1.06 | 3.69E-04 |
| TERTU_RS08990 | *ttpC3* | -0.81 | 2.55 | 0.03 |
| TERTU_RS08995 | *exbB3* | -0.84 | 1.82 | 0.03 |
| TERTU_RS09000 | *exbD3* | -0.6 | 1.65 | 0.16 |
| TERTU_RS09005 | *tonB3* | -1.12 | 2.54 | 2.20E-03 |
| TERTU_RS09010 | *ttpD3* | -0.6 | 4.11 | 0.07 |

**Table S4.** RNA-seq result to understand iron-regulation of genes potentially encoding TonB-dependent outer membrane receptor

| Locus tag | logFC | logCPM | FDR-adjusted p-value |
| --- | --- | --- | --- |
| TERTU_RS00400 | -0.50 | 5.33 | 0.16 |
| TERTU_RS02235 | -2.90 | 13.53 | 3.18E-08 |
| TERTU_RS02920 | 0.02 | 9.41 | 0.97 |
| TERTU_RS03510 | -0.02 | 6.33 | 0.97 |
| TERTU_RS05530 | -1.72 | 5.88 | 1.14E-08 |
| TERTU_RS05545 | 2.88 | 10.49 | 7.93E-13 |
| TERTU_RS05620 | -0.22 | 4.95 | 0.56 |
| TERTU_RS06590 | -0.85 | 5.08 | 0.01 |
| TERTU_RS06645 | -3.05 | 7.41 | 2.40E-14 |
| TERTU_RS06655 | 0.03 | 12.84 | 0.95 |
| TERTU_RS06660 | -1.03 | 6.60 | 1.06E-03 |
| TERTU_RS08920 | 0.27 | 5.13 | 0.49 |
| TERTU_RS08980 | -0.42 | 4.39 | 0.21 |
| TERTU_RS09115 | 1.65 | 7.14 | 6.99E-04 |
| TERTU_RS10285 | -1.59 | 9.61 | 1.45E-03 |
| TERTU_RS10395 | -0.26 | 6.34 | 0.47 |
| TERTU_RS11325 | 6.33 | 12.22 | 2.48E-44 |
| TERTU_RS11935 | -1.37 | 11.73 | 5.43E-04 |
| TERTU_RS12735 | -1.83 | 4.66 | 2.04E-05 |
| TERTU_RS12820 | -0.13 | 6.73 | 0.76 |
| TERTU_RS14810 | 0.13 | 10.38 | 0.79 |
| TERTU_RS14845 | 0.60 | 6.81 | 0.10 |
| TERTU_RS14930 | -0.15 | 7.46 | 0.72 |
| TERTU_RS15340 | -5.98 | 9.09 | 7.14E-19 |
| TERTU_RS15625 | -2.16 | 5.58 | 5.97E-06 |
| TERTU_RS16015 | -1.26 | 8.11 | 8.80E-04 |
| TERTU_RS16355 | 0.44 | 7.52 | 0.24 |
| TERTU_RS16735 | 1.66 | 8.63 | 1.45E-05 |
| TERTU_RS17025 | -1.76 | 8.63 | 2.25E-05 |
| TERTU_RS17140 | 3.44 | 8.38 | 1.94E-19 |
| TERTU_RS17895 | 4.09 | 9.96 | 2.41E-20 |
| TERTU_RS18025 | 8.55 | 12.09 | 2.94E-64 |
| TERTU_RS18165 | -2.09 | 5.48 | 7.17E-13 |
| TERTU_RS18190 | -1.39 | 4.11 | 1.30E-05 |
| TERTU_RS18495 | -2.11 | 7.59 | 9.63E-07 |
| TERTU_RS19320 | 0.40 | 5.09 | 0.23 |
| TERTU_RS20475 | 0.76 | 4.31 | 0.02 |
| TERTU_RS20690 | -0.72 | 12.24 | 0.09 |

**Figure S1.**
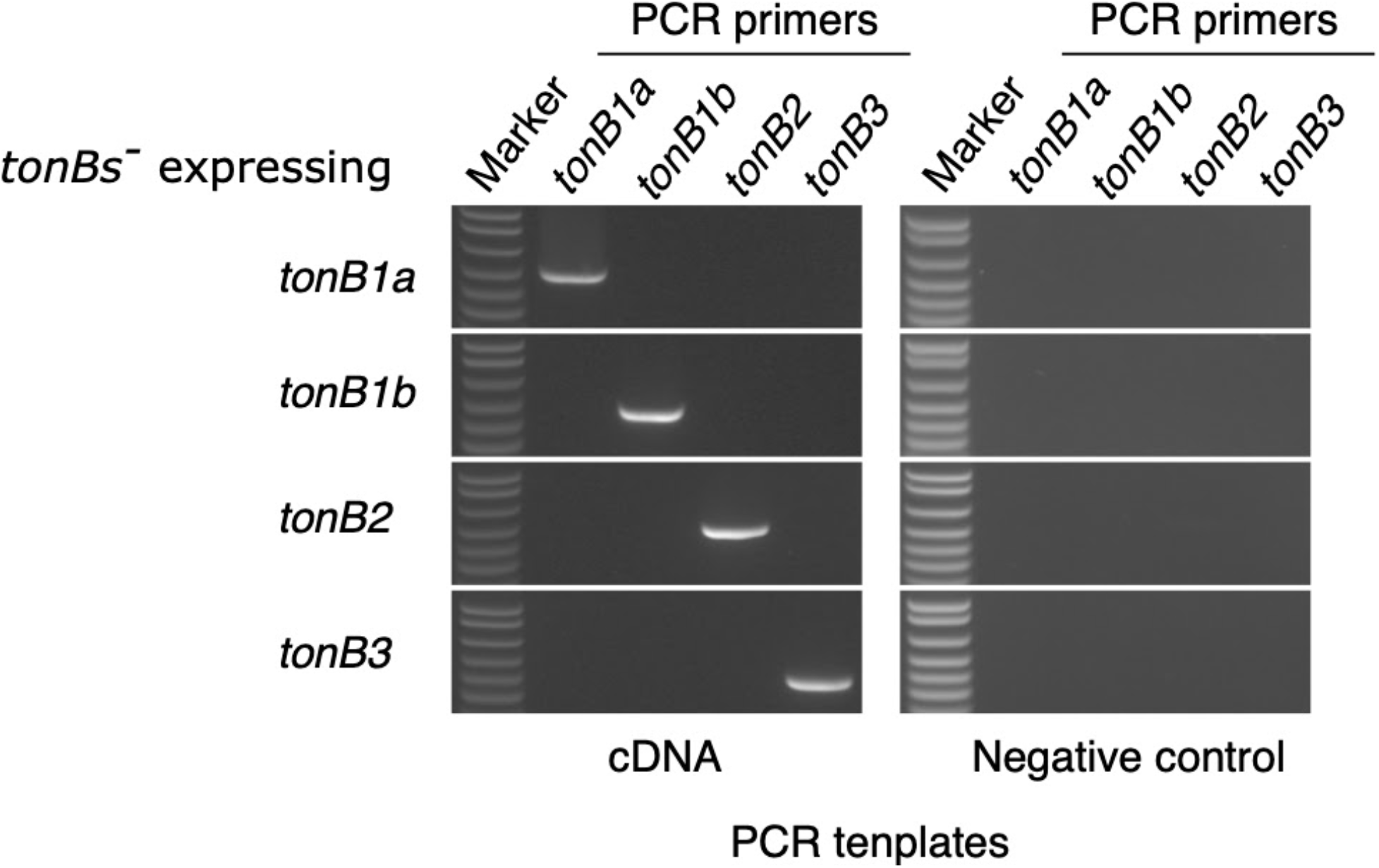
RT-PCR to confirm the expression of TonB genes. Each TonB gene was cloned into the expression vector pHN45 and conjugated into the *T. turnerae* T7901 *ΔtonB1abΔtonB2ΔtonB3* (tonBs-) strain in which all *tonB* genes were deleted. The strains were grown in SBM medium containing sucrose (0.5%), FAC (10 μM) and Km (50 μg/ml) at 30°C until reaching the exponential phase (OD_600_ 0.2-0.3). Negative control, RT-reaction without RT enzyme; FAC, ferric ammonium citrate; pHN45, plasmid expression vector.

## Fur titration assay

The fur titration assay (FURTA) was performed as described by (Stojiljkovic et al., 1994). DNA fragments to be tested were cloned into pBluescript II and transformed into E. coli H1717, and transformants were streaked on MacConkey agar plates supplemented with ammonium iron (II) sulfate (30μM) and Amp (100 μg/ml). Appearance of pink color (Lac+ phenotype, Fur-binding to cloned DNA fragments) around streaked *E. coli* was checked after overnight incubation at 37°C.

---

**Figure S2.**
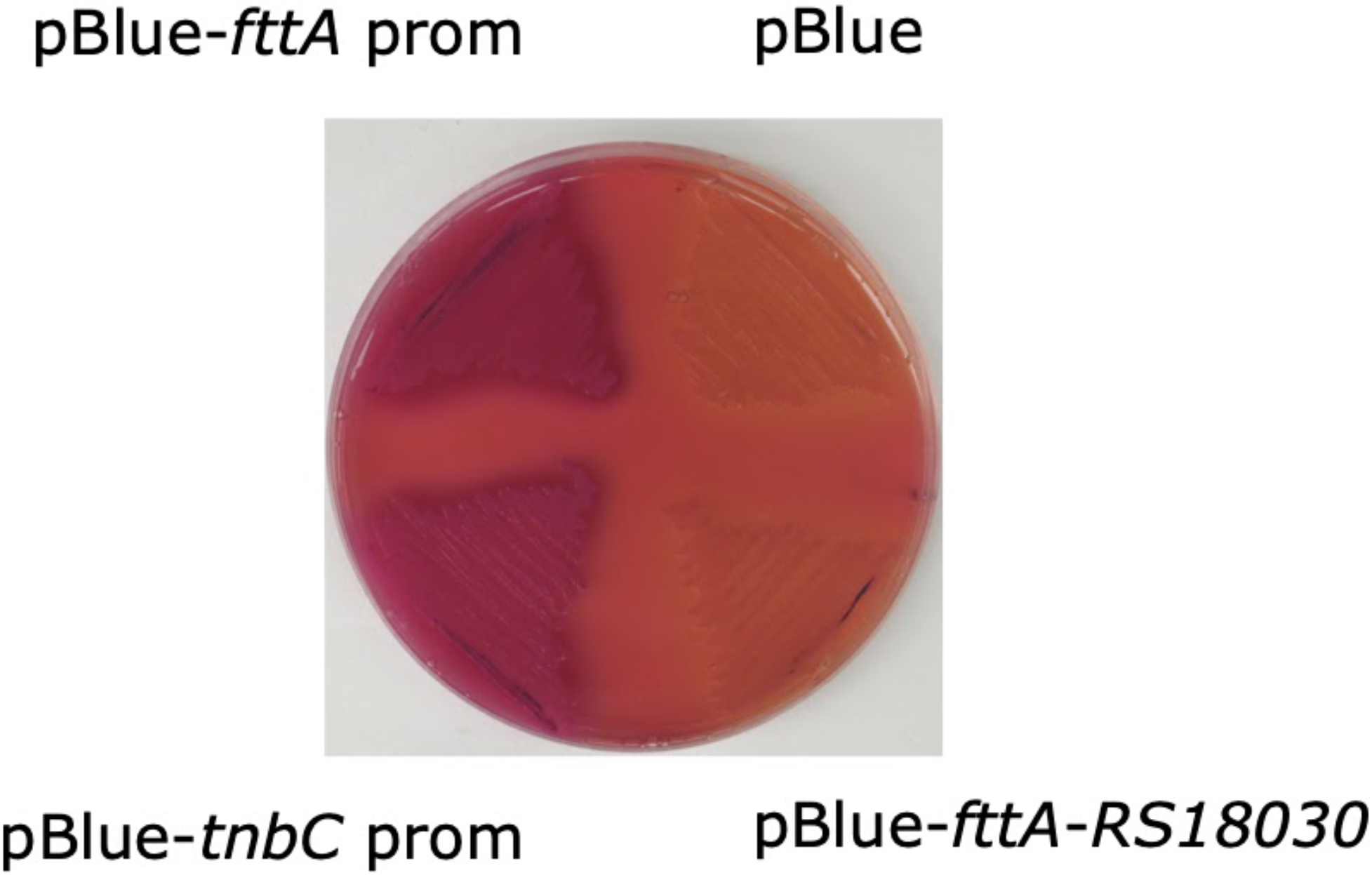
*E. coli* Fur can bind to the promoter regions of *fttA* and *tnbC*. Binding of *E. coli* Fur to promoter regions of *fttA, tnbC*, and the region between *fttA* and *TERTU_RS18030* (negative control) was tested by Fur titration assay (Stojiljkovic et al., 1994). *E. coli* H1717 containing plasmids harboring each fragment were streaked on MacConkey agar plates, and the presence of pink color (Lac+ phenotype) around streaked bacteria was evaluated after overnight incubation at 37°C. pBlue, pBluescript II; *fttA* prom, *fttA* promoter region; *tnbC* prom, *tnbC* promoter region; *fttA-RS18030*, the intergenic region between *fttA* and *TERTU_RS18030*.

## Notes

### Competing Interest Statement

The authors have declared no competing interest.

